# An oxygen-sensing *Polycomb*-group protein encodes flooding stress memory in plants

**DOI:** 10.64898/2026.05.01.722198

**Authors:** Aida Maric, Gabriele Meder, Carlos Rodriguez-Cisneros, Livia Schaub, Rory Osborne, Yixuan Fu, Sara Simonini, Daniel J. Gibbs, Sjon Hartman

**Affiliations:** Plant Environmental Signalling and Development, Faculty of Biology, University of Freiburg, 79104, Freiburg, Germany; CIBSS–Centre for Integrative Biological Signalling Studies, University of Freiburg, 79104, Freiburg, Germany; Spemann Graduate School of Biology and Medicine (SGBM), University of Freiburg, 79104, Freiburg, Germany; School of Biosciences, University of Birmingham, Edgbaston, B15 2TT, United Kingdom; Institute of Plant and Microbial Biology, University of Zurich, CH8008, Zurich, Switzerland

**Author notes:** Shared senior authors (Daniel J. Gibbs, Sjon Hartman).

## Abstract

Most organisms, including plants, can encode stress memories that improve resilience to repeated environmental challenges. Flooding events expose plants to recurrent hypoxia, yet whether plants establish an adaptive memory of flooding is unclear. Here we show that somatic flooding stress memory is a conserved feature across multiple angiosperm species. In *Arabidopsis*, this memory depends on the oxygen-sensitive *Polycomb* Repressive Complex 2 (PRC2) subunit VERNALIZATION2 (VRN2). Loss of VRN2 impairs epigenetic memory formation and disrupts transcriptional memory at key genes that promote anthocyanin accumulation and repress leaf senescence, adaptive responses that enhance flooding tolerance. Collectively, we reveal a molecular mechanism where VRN2-PRC2 acts as both an oxygen sensor and chromatin effector to establish adaptive flooding stress memory in plants.

**Short:** Our work defines a molecular mechanism for flooding stress memory in plants, actioned via the PRC2 subunit VRN2, that acts as the direct molecular (low) oxygen sensor and transducer. Coupled to its role in cold sensing, this identifies VRN2 as a master sensor subunit of the plant PRC2 that integrates multiple environmental cues to modulate the chromatin landscape and encode environmental memory.

## Introduction

Many organisms frequently encounter environmental stresses and can form epigenetic memories that shape their responses to subsequent stress (Harris et al., 2023; Houri-Zeevi et al., 2020). Such stress memory can provide organisms with enhanced tolerance to repeated environmental challenges and, in some cases, be transmitted across generations (Kim et al., 2025). In plants, somatic stress memory is a well-established phenomenon and has been studied primarily in the context of temperature and pathogen stress (Friedrich et al., 2021; Hannan Parker et al., 2022; Lämke & Bäurle, 2017). Flooding is a major abiotic stress that increasingly threatens plant productivity and causes substantial crop losses worldwide, yet little is known about whether plants can epigenetically encode memory of flooding stress (Gibbs et al., 2025; Maric et al., 2025; Rodriguez-Cisneros et al., 2026).

Plant submergence leads to oxygen deprivation (hypoxia), and the adaptive transcriptional response to hypoxia in flowering plants (angiosperms) is largely regulated by oxygen-dependent proteolysis of group VII Ethylene Response Factor (ERFVII) transcription factors via the PCO/N-degron pathway (Gibbs et al., 2011; Licausi et al., 2011). While oxygen sensing in plants has been primarily understood through transcription factor regulation, it remains unclear whether and how oxygen sensing can directly interface with chromatin-based gene regulation. Since the original discovery of oxygen sensing ERFVII proteins, only two additional oxygen-sensitive N-degron pathway substrates have been identified in plants (Gibbs et al., 2018; Weits et al., 2019), including the *Polycomb* Repressive Complex 2 (PRC2) subunit VERNALIZATION2 (VRN2), which is well positioned to connect oxygen sensing to the epigenetic control of gene expression.

PRC2 is a conserved chromatin-modifying complex that catalyses histone H3 lysine 27 trimethylation (H3K27me3) at target loci to mediate stable transcriptional repression and control diverse biological processes across eukaryotes (Mozgova & Hennig, 2015; Osborne, 2026). In *Arabidopsis thaliana* (*Arabidopsis*), PRC2 consists of four core subunits, including one of three interchangeable *Drosophila* Suppressor of Zeste 12 (Suz12) homologs, including VRN2. VRN2-PRC2 plays a key role during the vernalization response by mediating epigenetic memory of prolonged cold exposure, enabling vernalization-dependent angiosperms to flower after winter through stable repression of the floral repressor *FLOWERING LOCUS C* (*FLC*) (Bastow et al., 2004; Gendall et al., 2001). This established role in encoding environmental memory suggests that VRN2 may act more broadly as an integrator of environmental signals into stable chromatin states. VRN2 also accumulates in endogenous hypoxic niches within meristematic tissues and primordia, where VRN2-PRC2 tunes the epigenetic regulation of growth-related genes that drive plant development and leaf expansion (Labandera et al., 2021; Osborne et al., 2025).

Prolonged cold and submergence-induced hypoxia promote ectopic VRN2 stabilization outside of meristems, but the biological function of this broader VRN2 accumulation during flooding-induced hypoxia remains unclear (Gibbs et al., 2018). These post-translational dynamics suggest that oxygen-dependent stabilization of VRN2 may connect hypoxic stress perception to chromatin-based regulation. Here, we show that multiple angiosperm species establish an adaptive somatic memory of flooding stress. In *Arabidopsis*, this response depends on VRN2, and we further show that VRN2-PRC2-controlled transcriptional memory at tolerance-associated genes promotes anthocyanin accumulation and represses leaf senescence. Together, these findings show that the oxygen-sensitive PRC2 subunit VRN2 links hypoxia sensing to chromatin-based regulation, enabling the establishment of adaptive flooding stress memory in plants.

## Results

### Plants require VRN2 to encode flooding stress memory

To unravel if plants have a capacity to encode and benefit from flooding stress memory, *Arabidopsis* plants were primed with a mild submergence stress in the light and recovered for four days before they were subjected to more severe submergence stress in the dark (Fig. 1A; S1). Our results demonstrate that primed plants have strongly enhanced flooding stress survival and retain more biomass compared to naïve plants (Fig. 1B-D, S1), hereafter referred to as flooding stress memory. A priming duration of more than two days is required to promote tolerance to recurrent submergence stress (Fig. S2A-B), and the priming benefit is independent of developmental age and whether priming caused a growth penalty (Fig. S1). Moreover, we observed that *Arabidopsis* is able to benefit from a primed state for at least 14 days (Fig. S2C-D). Additionally, submergence priming promoted subsequent flooding tolerance in the wetland species *Rumex palustris* and cereal *Hordeum vulgare* (barley) (Fig. S3), revealing a potential conserved capacity for encoding flood stress memory in angiosperms. We next aimed to unravel how the model species *Arabidopsis* encodes flooding stress memory and functionally benefits from a primed state.

**Figure 1.**
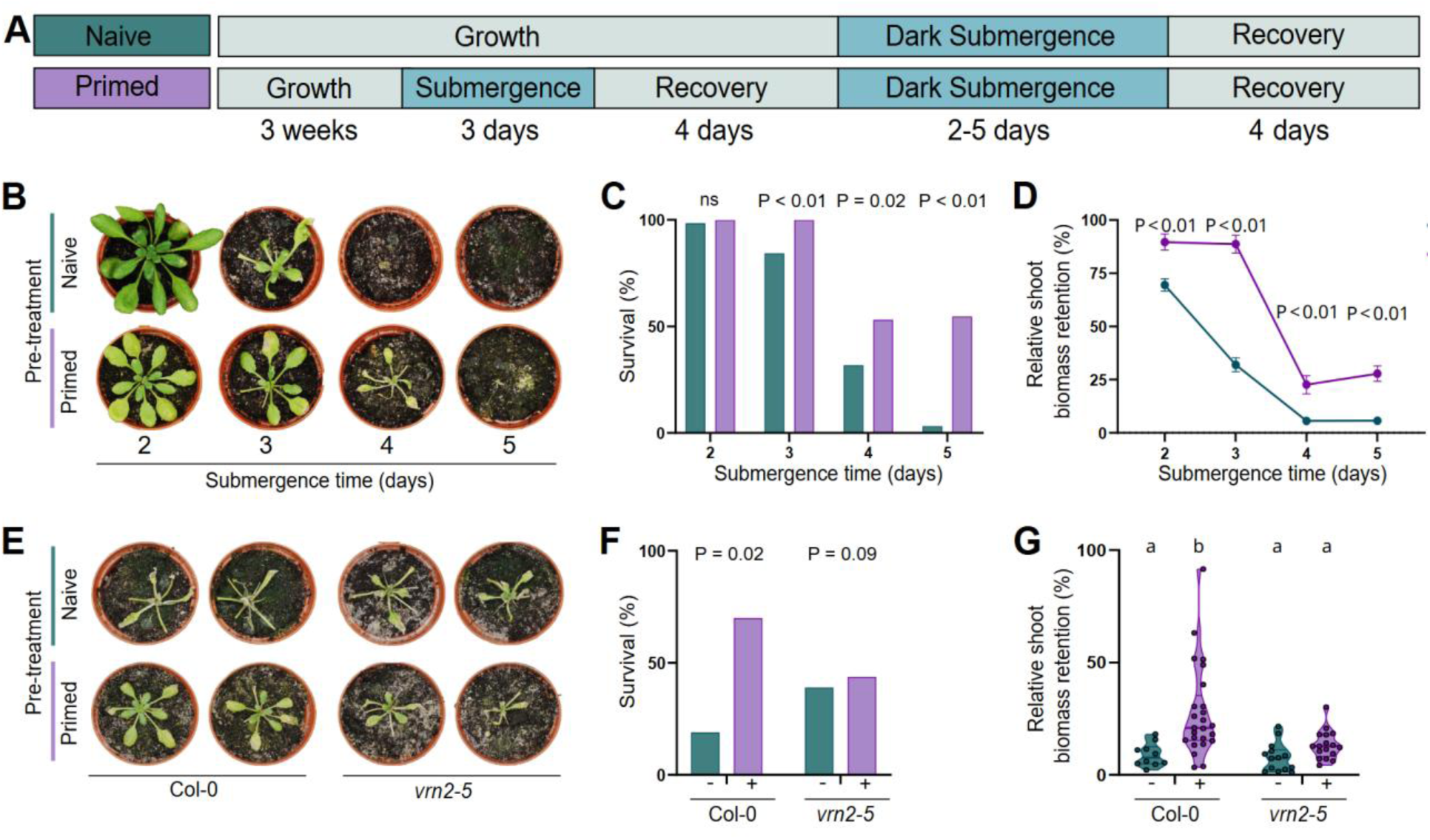
Submergence priming promotes VRN2-dependent tolerance to recurring flooding stress. **(A)** Experimental setup to quantify the effects of light submergence (priming) treatments on subsequent dark submergence tolerance. **(B)** Representative images, **(C)** plant survival and **(D)** relative shoot biomass retention of naïve (teal) and submergence primed (violet) Arabidopsis plants after recovery from 2-5 days of dark submergence. **(E)** Representative images, **(F)** plant survival and **(G)** relative shoot biomass retention of naïve (teal) and submergence primed (violet) Arabidopsis wild type Col-0 and *vrn2-5* mutant plants after recovery from 4 days of dark submergence. Data shown in C & F represent the mean values of two experimental replicates; D & G show data of surviving plants from these two respective experimental replicates (10 ≤ n ≥ 32). Statistically significant differences were calculated by Fisher’s exact test (C & F), two-tailed Student’s t test (D), and two-way ANOVA with Tukey’s HSD (P < 0.05; G).

The PRC2 subunit VRN2 is a direct biochemical sensor of low oxygen that is enriched in plant meristems, but accumulates more broadly in response to acute hypoxia and submergence, identifying it as a potential mediator of hypoxia-responsive epigenetic regulation (Gibbs et al., 2018; Labandera et al., 2021; Osborne et al., 2025). Since flooding imposes hypoxia, we tested whether *Arabidopsis* plants lacking VRN2 could still remember flooding stress. Strikingly, *vrn2-5* loss-of-function plants are unable to benefit from a primed state to overcome sequential flooding stress (Fig. 1E-G). Complementation of *vrn2-5* with a *pVRN2::VRN2-YFP* construct (Osborne et al., 2025), restored the ability to benefit from submergence priming (Fig. S4). Importantly, basal flooding tolerance is not impaired in *vrn2-5* plants. Collectively, these results reveal that VRN2 is essential for flooding stress memory in *Arabidopsis*.

### Submergence priming leads to chromatin remodeling and two modes of transcriptional memory

“Transcriptional memory” describes altered gene expression behavior in primed compared to naïve plants, may contribute to memory-related phenotypes, and is strongly dependent on epigenetic mechanisms (Lämke & Bäurle, 2017). Transcriptional memory can be classified as either a long-lasting sustained transcriptional state after a priming stimulus (type I) or a modified transcriptional activation pattern upon a recurring stress (type II) (Oberkofler et al., 2021). We performed RNA sequencing to quantify these two modes of transcriptional memory in primed compared to naïve *Arabidopsis* rosettes (Fig. 2A). Because the flooding memory effect is VRN2-dependent (Fig. 1E-G, S4), we focused specifically on the subset of differentially expressed genes that occurred exclusively in primed compared to naïve wild-type plants, but not the *vrn2-5* mutant. Primed wild-type plants maintained a much larger sustained (type I) transcriptional state compared to *vrn2-5* (Fig. 2B, Data S1). Gene Ontology (GO) term enrichment of type I genes specific to wild type revealed an overrepresentation of terms related to anthocyanin biosynthesis and response to light in the upregulated gene set, and biotic stress and proteostasis in the downregulated gene set (Fig. 2C, Data S2). During subsequent dark submergence stress, primed plants of both wild-type and *vrn2-5* generally showed a milder type II transcriptional response (when compared to type I, Fig. 2D, Data S1). Here, upregulated wild type-exclusive type II GO terms were associated with iron signaling, lignin metabolism and flavone biosynthesis, whereas the downregulated type II genes were enriched for diverse GO term processes that include the regulation of leaf senescence (Fig. 2E, Data S2), which has previously been shown to attenuate flooding stress tolerance (Fukao et al., 2012; Rankenberg et al., 2024; Yeung et al., 2018).

**Figure 2.**
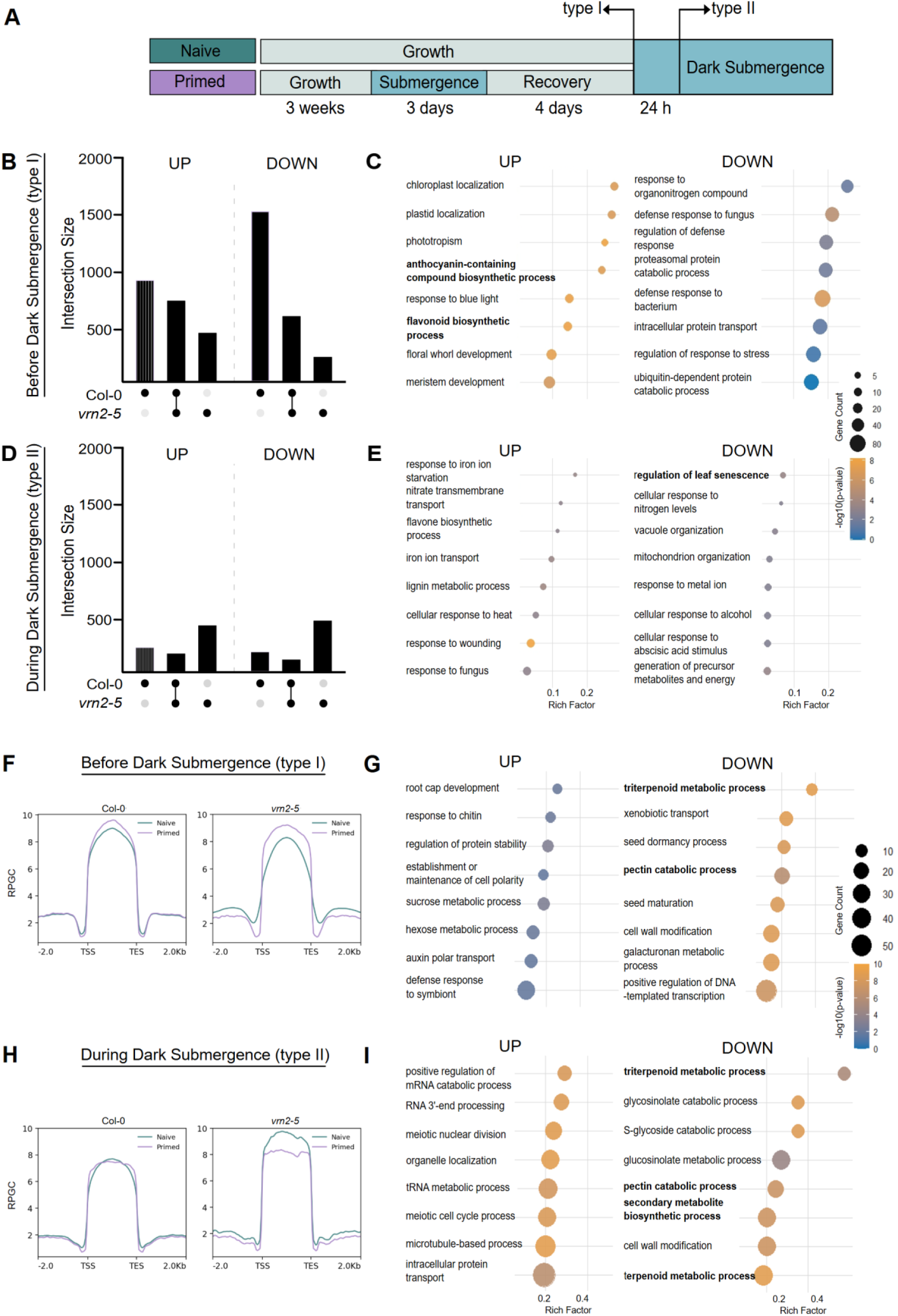
Submergence priming induces two modes of transcriptional memory and chromatin remodeling. **(A)** Experimental setup indicating sampling timepoints to quantify differential genome-wide transcript and H3K27me3 levels in naïve and primed plants before (type I) and during (type II) the triggering submergence stress treatment. **(B, D)** UpSet plot showing the number of DEGs (primed vs naïve) exclusive to Col-0 wild type and *vrn2-5* primed plants, and in both genotypes, before (B) and after (D) 24h of dark submergence. The up- and downregulated genes are shown in left and right, respectively. **(C, E)** GO enrichment analysis of genes differentially expressed exclusively in primed Col-0 wildtype plants **(F, H)** Metagene plot showing H3K27me3 / H3 levels in primed vs naïve wildtype and *vrn2-5* plants before (F) and after (H) 24h of dark submergence stress. RPGC; reads per genomic content; TSS, transcription start site; TTS, transcription termination site. **(G, I)** GO term enrichment analysis for genes that exclusively gained or lost H3K27me3 levels in primed wild-type plants before (G) and after (I) 24h of dark submergence stress.

VRN2-PRC2 canonically acts through depositing H3K27me3 to repress gene expression. H3K27me3 is often mitotically stable and its deposition on target loci can thus act as a memory mark to modify transcriptional activity (Bastow et al., 2004; Coleman & Struhl, 2017; Osborne et al., 2025; Zenk et al., 2017). Accordingly, we quantified the VRN2-dependent genome-wide H3K27me3 landscape in primed and naïve plants using the enzyme-tethering epigenome profiling technique CUT&Tag (Fu et al., 2026). Before the triggering stress, primed plants showed slightly enhanced H3K27me3 enrichment in gene bodies compared to naïve plants (Fig. 2F, S5, Data S3). Interestingly, whilst *vrn2-5* plants showed slightly reduced H3K27me3 levels compared to wildtype, the priming-induced H3K27me3 enrichment was still evident in *vrn2-5* plants. Upon the triggering submergence stress, no global H3K27me3 differences were observed between naïve and primed wildtype plants, but naïve *vrn2-5* plants showed slight global H3K27me3 enrichment (Fig. 2H, S5, Data S3). These results suggest that submergence priming generally triggers a VRN2-independent global H32K27me3 increase, reminiscent of what occurs during in heat stress priming (Yamaguchi et al., 2021), potentially leading to general transcriptional repression (Fig. 2B, Data S1, S3). In addition, the results indicate that especially during submergence, VRN2-PRC2 likely targets a specific subset of genes, rather than extensive chromatin-based remodeling of the genome. Accordingly, we investigated which genes and processes are overrepresented in H3K27me3 enrichment profiles. Interestingly, priming led to H3K27me3 depletion at loci associated with triterpenoid and secondary metabolite biosynthesis, and lignin and pectin catabolism (Fig. 2G, I, Data S4), potentially explaining its corresponding transcriptional activation profile in primed plants (Fig. 2C, Fig. S6). VRN2-dependent (wild-type exclusive) loci showing H3K27me3 enrichment in primed plants were associated with processes that include sugar signalling, auxin transport but also senescence-related hormonal responses (Fig. 2G, I, Data S4). Collectively, these results suggest that (1) priming leads to a global reduction of H3K27me3 deposition in gene bodies, and (2) VRN2-PRC2-dependent H3K27me3 deposition is highly specific to a subset of target loci.

### VRN2- and CHS-dependent anthocyanin biosynthesis contribute to flooding stress tolerance

The identification of pigment-like secondary metabolites, flavonoid- and anthocyanin-associated processes in our combined RNA-seq and chromatin analyses (Fig. 2C, G, I, S6, Data S1-4) prompted us to investigate whether priming triggers anthocyanin production and contributes to flooding stress memory. Plants use secondary metabolites to cope with a wide range of adverse environmental conditions. Among these, flavonoids and their anthocyanin derivatives, were shown to have protective roles through antioxidant capacity, functioning as signaling molecules, or mitigation of high light, drought and salt stress (Davies et al., 2018). Moreover, high flavonoid and anthocyanin content correlate with improved flooding tolerance of quinoa plants, but functional and causal evidence is still lacking (Jiang et al., 2025; Yu et al., 2023). We found that all genes encoding for the enzymes catalyzing anthocyanin biosynthesis, including *TRANSPARENT TESTA 4* (*TT4*) which encodes the chalcone synthase (CHS) enzyme that catalyzes the initial step of flavonoid biosynthesis (Peer et al., 2001), showed sustained type I transcriptional memory in a VRN2-dependent manner (Fig. 3A-C. S6). Interestingly, this type 1 transcriptional memory pattern correlated with strongly reduced H3K27me3 levels for *TT4 and TT3,* and partially reduced H3K27me3 for *TT7* and *TT5* in primed plants (Fig. 3D, Fig. S6), indicating that they are not likely directly targeted by VRN2-PRC2, but that their priming-induced demethylation still depends on its activity.

**Figure 3.**
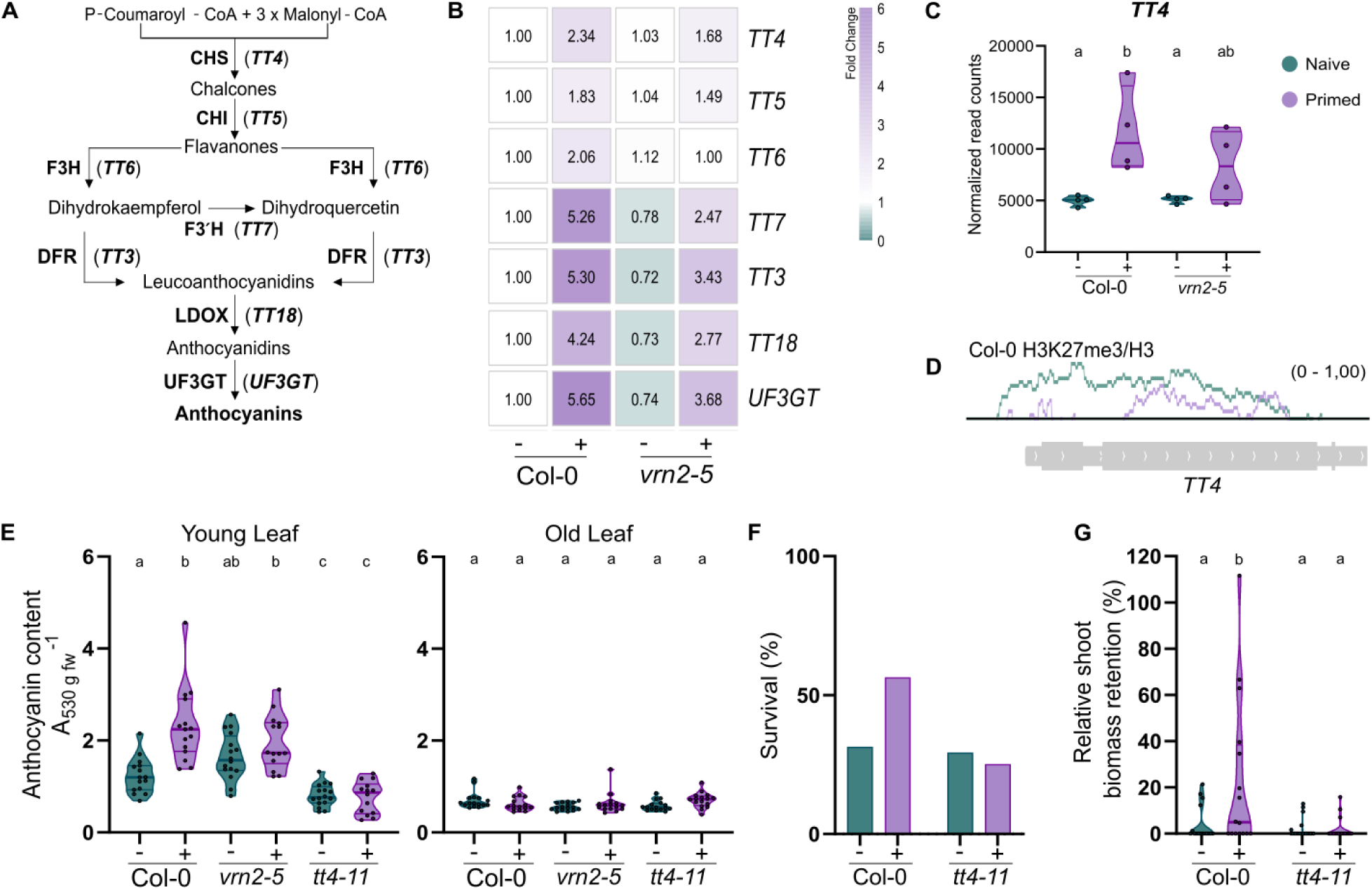
Submergence priming triggers VRN2-mediated anthocyanin accumulation promoting flooding tolerance. **(A)** Overview of the Arabidopsis anthocyanin biosynthesis pathway, including key enzymes that catalyze specific steps (and their encoding genes), based on (Peer et al., 2001). **(B)** Sustained type I transcriptional memory of key anthocyanin biosynthesis-related genes, and **(C)** *TT4* mRNA read count and **(D)** *TT4* locus coverage plot of H3K27me3 relative to H3 in the indicated naïve (- / teal) and submergence primed (+ / violet) rosettes, before the triggering submergence treatment. Values are presented relative to naïve Col-0 samples. **(E)** Anthocyanin content of primed (+) and naïve (-) Col-0, *vrn2-5* and *tt4-11* young (left) and old (right) leaf samples before the triggering submergence treatment. **(F)** Survival and **(G)** relative rosette fresh weight of primed (+) and naïve (-) Col-0 and *tt4-11* plants after recovery of 4-day dark submergence compared to dark controls (Fig. S1). In E & G the results from two independent experimental repeats are shown (n ≥ 7 per experiment). Letters in C, E & G indicate statistically similar groups determined by two-way ANOVA with Tukey’s HSD (P < 0.05).

To test if this type I transcriptional memory pattern of anthocyanin biosynthesis genes (Fig. 3A-C) translates into changes in anthocyanin production, we assessed the anthocyanin levels in naïve and primed plants of wild type, *vrn2-5* and *tt4-11* (Fig. 3E). As previous research showed submergence- and hypoxia-induced accumulation of VRN2 specifically in younger leaves (Gibbs et al., 2018; Labandera et al., 2021; Osborne et al., 2025), and because we observed that younger leaves typically survive flooding stress in primed plants (Fig. 1), we distinguished between younger and older tissues for our analysis. Young leaves of primed wild-type plants accumulated more anthocyanins compared to their naïve counterparts, and this difference was attenuated in *vrn2-5* mutant plants, correlating with respective gene expression patterns (Fig. 3E). As expected, the *tt4-11* loss-of-function mutant showed reduced anthocyanin levels in both treatments, and no differences in anthocyanin content were observed in the older leaves of any of the genotypes. Because priming promotes VRN2-dependent anthocyanin accumulation, we further assessed the ability of *tt4-11* mutants lacking anthocyanin accumulation to benefit from submergence priming. Strikingly, we found that *tt4-11* plants no longer benefited from priming and generally were slightly more sensitive to flooding stress (Fig. 3F, G). These results reveal that VRN2-dependent sustained transcriptional memory of CHS-controlled pigmentation in primed plants contributes to flooding stress memory in Arabidopsis.

### VRN2-dependent priming attenuates leaf senescence during dark submergence

Our omics analysis identified senescence-related genes as a putative VRN2-dependent targets, indicating that regulation of this process could contribute to flooding stress memory (Fig. 2E). Leaf senescence is the tightly controlled final stage of leaf development that ensures nutrient allocation to younger tissues (Lim et al., 2007). Senescence is preferentially induced in older leaves, and PRC2-mediated H3K27me3 deposition and REF6-mediated H3K27me3 demethylation occur at key senescence regulatory gene loci, and repress and promote leaf senescence, respectively (Liu et al., 2019; Wang et al., 2019). Genetic interference of leaf senescence was previously shown to promote dark submergence and re-oxygenation resilience in *Arabidopsis* and rice (Fukao et al., 2012; Rankenberg et al., 2024; Yeung et al., 2018). We observed a VRN2-dependent enrichment of leaf senescence-related genes that showed type II memory in primed plants as part of the transcriptional response to dark submergence stress (Fig. 2E, S7), characterized by reduced induction of genes including key senescence regulators *ANAC055* and *Non-Yellowing Color 1* (*NYC1*) (Fig. 4A-C, Fig. S7). We further observed that some of these loci, including *ANAC055* but not *NYC1*, showed VRN2-dependent enrichment of the repressive H3K27me3 mark its promoter region before the triggering dark submergence stress (Fig. 4A-C, S8, Data S4), implicating *NAC055* in a potential direct VRN2-PRC2-targeted mechanism for the observed attenuated transcriptional memory of senescence genes.

**Figure 4.**
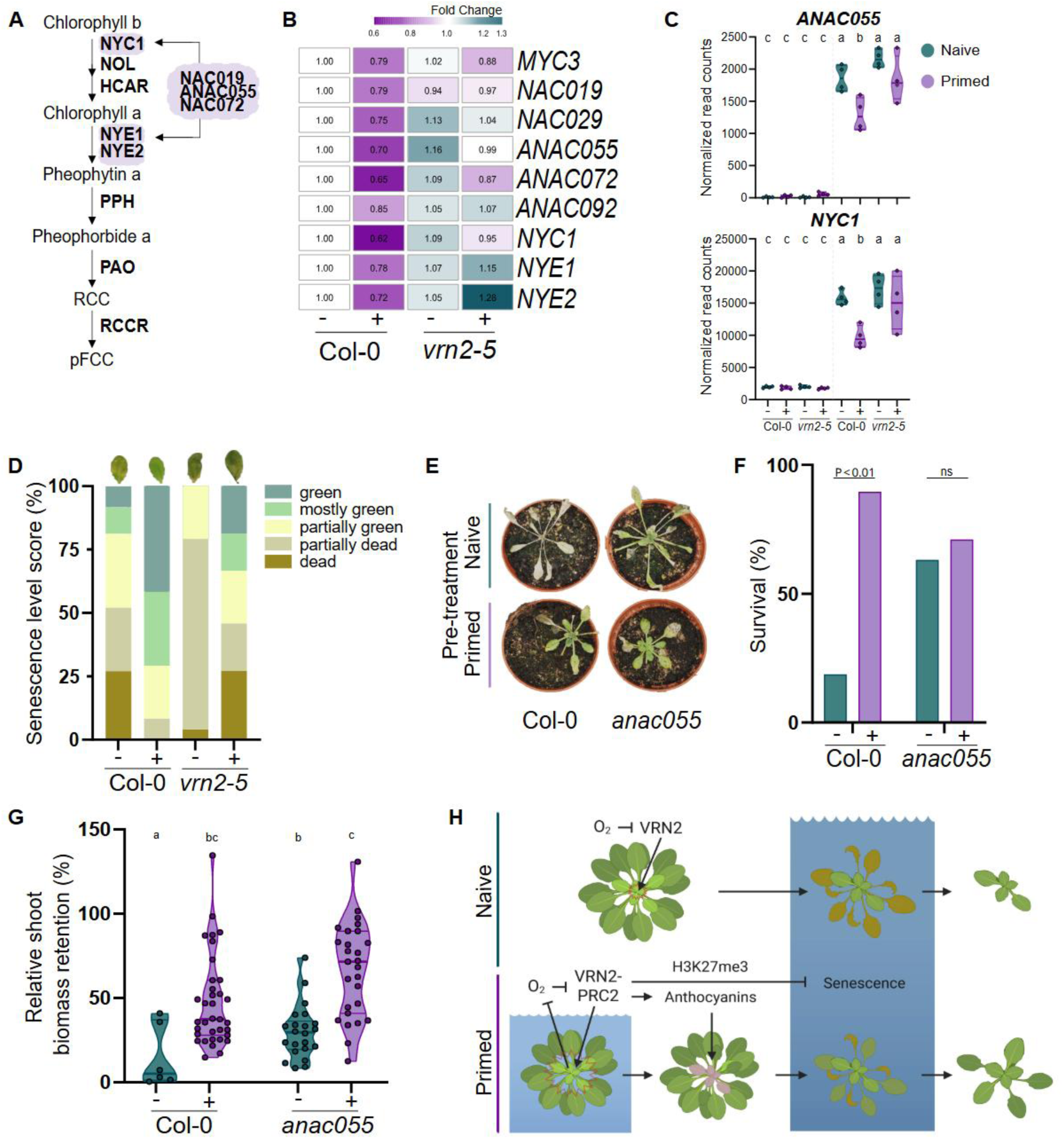
Priming represses senescence and *ANAC055* in *Arabidopsis* rosettes, promoting flooding stress tolerance. **(A)** Selected overview of the Arabidopsis leaf senescence and chlorophyll degradation pathway, highlighting (including key regulatory transcription factor) genes that show reduced transcriptional activation after priming. **(B)** Relative mRNA abundance of senescence associated genes in primed/naïve wild-type and *vrn2-5* plants during dark submergence. **(C)** *ANAC055* and *NYC1* transcriptional memory patterns before and during dark submergence. **(D)** Young rosette leaves (youngest leaf 3 to 5) were scored according to the senescence levels in primed (+) and naïve (-) plants after dark-induced senescence, from 1 (dead) to 5 (fully green). The ratio of each category was expressed as the percentage of total leaves considered (n = 12-16). **(E)** Representative images, **(F)** plant survival and **(G)** relative shoot biomass retention of naïve (teal) and primed (violet) Arabidopsis wild type Col-0 and *anac055* mutant plants after 4 days of recovery from dark submergence. Letters in C and G indicate statistically similar groups determined by two-way ANOVA with Tukey’s HSD (P < 0.05). P-values in F are result of Fisher’s exact test. Data come from at least three independent experimental replicates (n > 8). **(H)** Proposed molecular mechanism of VRN2-controlled flooding stress memory in Arabidopsis. In naïve plants, O_2_-controlled VRN2 is restricted to the hypoxic shoot apical meristem and plants under dark submergence have high leaf senescence, limiting stress recovery. In primed plants, VRN2 accumulates in younger leaves and VRN2-PRC2 deposits H3K27me3 marks in chromatin, enhancing anthocyanin biosynthesis during recovery and limiting leaf senescence upon sequential submergence stress, promoting flooding tolerance.

Therefore, we next assessed if primed plants promote flooding resilience by limiting chlorosis and *NAC055* expression in response to dark-induced senescence. We observed that young wild-type leaves remained greener than the corresponding naïve controls, and that this effect was reduced in *vrn2-5* plants (Fig. 4D-E). Interestingly, in addition to its role in senescence, *ANAC055* was shown to be negative regulator of heat stress memory in *Arabidopsis* seedlings (Alshareef et al., 2022). We thus tested whether ANAC055 is also involved in flooding stress memory and found that *anac055* loss-of-function mutants showed significantly enhanced flooding tolerance relative to wild type and only mildly benefited from priming (Fig. 4E). Collectively, these results suggest that ANAC055 negatively regulates flooding stress tolerance, and that its VRN2-dependent suppression can contribute to memory-mediated tolerance in part by limiting flooding-induced senescence.

## Discussion

Our work demonstrates that plants can encode somatic flooding stress memory to promote resilience against recurring stress (Fig. 1), and that this phenomenon is conserved in several angiosperm species (Fig. S3). Strikingly, oxygen-sensitive proteolytic control of VRN2 arose early during angiosperm evolution and is present in *Arabidopsis*, barley and rice (Gibbs et al., 2018), suggesting that VRN2-dependent flooding memory may be a distinct feature of angiosperms. Our findings show that dynamic H3K27me3 methylation, alongside oxygen-sensitive histone lysine demethylation (Chakraborty et al., 2019; D. Zhang et al., 2025), plays a role in oxygen-dependent chromatin reprogramming, and reveals a mechanism underlying PRC2-mediated hypoxia stress memory (Kim et al., 2025).

Moreover, we demonstrate that VRN2-PRC2 mediates H3K27me3 deposition on specific memory loci and controls two modes of transcriptional memory that functionally contribute to flooding stress memory, including sustained *TT4*-mediated anthocyanin accumulation and the repressed induction of senescence in younger leaves during dark submergence (Fig. 4F). Endogenous VRN2 accumulation in plant meristems spreads to younger leaves under flooding-induced hypoxia (Gibbs et al., 2018), and may thus specifically mark and protect these younger leaves during submergence priming for sequential flooding stress protection (Fig. 4F). Whilst anthocyanin accumulation and reduced senescence in primed plants are at least partially VRN2-dependent (Fig. 2-4), it remains to be investigated which loci are directly targeted by VRN2-PRC2 for H3K27me3 deposition (Fig. S6, 8), and to what extend this putative memory mark controls transcriptional behavior. In the future, the use of epigenome editing tools could aid in demonstrating such a causal relationship between locus-specific memory marking and transcriptional activity (Policarpi et al., 2024), in addition to controlling transcriptional memory of tolerance genes such as *TT4* and *ANAC055* to promote flooding resilience without the requirement of a memory-encoding priming treatment. In conclusion, we identify a novel mechanism of flooding resilience that emerges specifically under repeated submergence stress, which could be (epi)genetically targeted to enhance flooding tolerance and stress memory.

## Acknowledgements

We thank Sabine Kenz, Marita Hermann, Lena Scheibe, Constanze Mohr, Jürgen Schmidt and Dirk Renz for technical support and the Plant Environmental Signalling and Development group, Thomas Laux and Frauke Garbsch of the Faculty of Biology, University Freiburg, for fruitful discussions.

## Author contributions

Conceptualization: A.M., D.J.G., S.H.; Methodology: A.M., R.O., Y.F., S.S., D.J.G., S.H.; Investigation: A.M., G.M., C.R.C., L.S., S.H.; Funding acquisition: A.M., D.J.G., S.H.; Supervision: A.M., D.J.G., S.H.; Writing – original draft: A.M., S.H.; Writing – review & editing: A.M., S.S. D.J.G., S.H.

## Data and materials availability

No restrictions are placed on materials, such as materials transfer agreements. All data are available in the main text or the supplementary materials. Raw sequencing data will be made available on a public database upon acceptance of the manuscript.

## Supplemental Files

Supplementary figures S1-S8

Supplementary data files S1-S4

## Materials & Methods

### Plant material and growth conditions

All seeds were grown on potting soil/sand mix (1:1) in a walk-in growth chamber under neutral-day length conditions (12h light at 21°C, 12h dark at 20°C; PAR: 100 μmol m^−2^s^−1^) and 70% relative humidity. Seeds were stratified for at least three days in the dark at 4°C to promote homogeneous germination before being transferred to the growth chamber. *Arabidopsis thaliana* of ecotype Columbia (Col-0) was used as the wild-type background for all the mutant lines in this study. The *vrn2-5* (SALK_201153) mutant and *vrn2-5*/pVRN2::VRN2-YFP transgenic line were described previously (Gibbs et al., 2018; Osborne et al., 2025), as well as *Rumex palustris* and *Hordeum vulgare* seeds (Hartman et al., 2019). Other mutant lines were kindly provided as follows: *tt4-11* (SALK_020583) by Prof. Brenda S.J. Winkel, Virginia Tech, USA (Bowerman et al., 2012) and *anac055* (SALK_011069) by Prof. Salma Balazadeh, Georg-August-Universität Göttingen, Germany (Alshareef et al., 2022).

### Flooding stress memory experiments

For the flooding priming and subsequent stress tolerance treatments (Fig. S1), three weeks old quasi-randomized plants were divided in two groups. One set of plants was light-submerged (primed) for three days while the other set (control) was submerged shortly (approx. ∼1 minute, to mimic soil water content and compaction changes of the submerged plants) and left to grow normally. The submergence always started four hours after the lights turned on (ZT4). After three days of priming submergence, the treated set of plants was de-submerged and left to grow normally with control plants for the next four days. Afterwards, both the control and primed sets of plants were further divided in half. One half of control plants and one half of primed plants were dark-submerged. The remaining control and primed plants were used as controls for naïve- and primed-submerged plants, respectively. In experiments where additional mutant genotypes were included, they were equally distributed and combined within the same tray of wild-type plants to remove potential growth tray and submergence tray effects. After the indicated dark submergence timepoints (indicated in figure legends, typically four days in main figures), plants were removed from water and left to recover for four more days. The plants were scored for survival and they were considered alive if the shoot apical meristem was viable (presence of green tissue). Shoot fresh weight was quantified by weighing only the green tissue per surviving plant. The relative shoot biomass retention was calculated as percentage mass of each treated plant relative to the respective pre-treatment (naïve/primed) control plantś mean biomass (see Fig. S1).

### Leaf Senescence Analysis

Experiments for leaf senescence analysis were performed similar to the previously described memory experiments, with some modifications. Briefly, after the 3-day priming stress, and 4-day recovery, plants were moved to dark to trigger dark-induced senescence. After 10 days of darkness, rosette leaves were detached, photographed and scored for senescence. Leaves considered as ‘old’ are the first true leaves; while leaves considered as ‘new’ are the last ones with a visible petiole. Senescence categories were assigned based on previously published research, with some modifications (Rankenberg et al., 2024). Briefly, leaf senescence scores were assigned as ‘green’ (fully green), ‘mostly green’ (∼10% - 30% yellow surface), ‘partially green’ (∼50% yellow surface), ‘partially dead’ (partially dead leaves) and ‘dead’ (completely dead leaves). Senescence scores were expressed as the ratio of leaves belonging to each group over total number of leaves observed of a specific leaf age.

### Anthocyanin measurements

For anthocyanin measurements, the young and old leaves were collected separately. Leaves considered as ‘old’ are the first true leaves; while leaves considered as ‘new’ are the last ones with a visible petiole. Anthocyanin measurements were performed as previously published, with modifications (Dieckmann et al., 2025; Jeong et al., 2010). Shortly, detached leaves were weighed, flash-frozen, ground into powder and mixed with 600 μL of extraction buffer [1% HCl in methanol (v/v)]. The samples were then incubated on a shaker at 4°C overnight. Extract was then thoroughly mixed with 800 μL of water and chloroform (1:1). Samples were the centrifuged at 12 000 rpm for 2 min and the moved to a multi-well plate. Absorbance was measured at 530, 657 and 720 nm using a plate reader (TECAN Infinite® 200 PRO) and anthocyanin content calculated using the following formula: [(A530–A720)–0.25×(A657–A720)] FW^−1^ (Dieckmann et al., 2025; Jeong et al., 2010).

### RNA-sequencing and bioinformatic analysis

Plants were grown for stress memory experiments and two whole rosettes were collected per sample by snap-freezing in liquid nitrogen at indicated timepoints (Fig. 2A). Four biological replicates were used per timepoint, treatment and genotype. Total RNA was extracted using the QIAGEN RNeasy® Plant Mini Kit and used for subsequent sequencing performed by Novogene Co., Ltd. Sequencing libraries were prepared using polyA enrichment method and sequencing was done on a NovaSeq X Plus platform, using a paired-end 150 bp per read strategy.

RNA sequencing data were analyzed using Galaxy platform and its public server at *usegalaxy.eu* (Abueg et al., 2024). Raw reads were cleaned up and trimmed using Cutadapt (Martin, 2011). Specifically, we removed the adapters, any reads with quality cutoff less than 20, and any reads shorter than 20 bp. Quality of cleaned up data was checked and summarized using MultiQC (Ewels et al., 2016). High-quality reads were then mapped to TAIR10 genome using RNA STAR mapper (Dobin et al., 2013) and counted with featureCounts (Liao et al., 2014). Differentially expressed genes were identified with DESeq2 (q-value < 0.05) (Love et al., 2014). Downstream analysis and visualization were performed using RStudio version 4.4.0 (https://www.r-project.org/) and GraphPad Prism version 8 (https://www.graphpad.com). Gene ontology (GO) enrichment analysis was performed using R package clusterProfiler with *org.At.tair.db* annotation (G. Yu et al., 2012). Plots for heatmaps and gene ontology data were generated using R packages *pheatmap* and *ggplot2 (Raivo Kolde, 2025; Wickham, 2016)*.

### CUT&Tag sequencing

Plants were grown for stress memory experiments and two whole rosettes were collected per sample by snap-freezing in liquid nitrogen at indicated timepoints (Fig. 2A). Two biological replicates were used per timepoint, treatment and genotype. CUT&Tag was performed as described before (Fu et al., 2026; Kaya-Okur et al., 2019), with some modifications. Specifically, tissues were ground and collected in DNA LowBind microtubes (Sarstedt). 1 ml of Honda buffer was added to each sample and nuclei were isolated by incubating on ice for 20 minutes, with occasional stirring. Samples were passed through 40 μm pluriStrainer (pluriSelect) and centrifuged for 15 minutes at 4000 x g and 4°C. The supernatant was discarded and 1 mL of LB01 buffer was added to the pellet. Samples were centrifuged again for 15 minutes at 1000 x g and 4°C. Resulting supernatant was discarded and pellet resuspended in 200 ul of Wash buffer. Concavalin-A beads (BioMag Plus Concanavalin A, Polyscience Inc, #86057-10) were activated in Binding buffer and 15 ul of activated beads were added to each sample. The mix was incubated on rotor for 10 minutes. The tubes were then placed on a magnetic stand (Invitrogen #CS15000) and the supernatant discarded. The sample was immediately resuspended in 50 μL of H3K27me3 (Cell signaling technology #9733408) and H3 (Active Motif #39064) antibodies, diluted in Antibody buffer at a 1:50 and 1:150 ratio, respectively. After the overnight incubation at 4°C, the samples were placed on a magnetic stand and supernatant removed. The sample was then immediately resuspended in 60 μL of secondary antibodies (Guinea Pig anti-Rabbit IgG: Antibodies-online #ABIN101961412; and Rabbit anti-Mouse IgG: Abcam #46540) diluted in Tween-Wash Buffer according to manufacturers’ instructions. After a 45-minute incubation at room temperature with secondary antibody, samples were placed on a magnetic stand and supernatant was removed. The beads were washed once with 500 μL Tween-wash buffer and buffer was completely removed. The samples were then resuspended in 60 μL of pA-Tn5 (produced in-house, loaded with adaptors and checked for activity) diluted in Tween-300-Wash buffer. After a 1h incubation, the beads were washed twice with 500 μL and 200 μL of Tween-300-wash Buffer, respectively. The tagmentation was then done by adding 300 μL of Tagmentation Buffer and incubation at 37°C for 1 hour. Tagmentation was then blocked by adding 10 μL of 0.5M EDTA (Roth #8040), 3 μL of 10% SDS (Invitrogen #AM9822), and 2.5 μL 20 mg/mL Proteinase K (Thermo Scientific #EO0492) and incubation at 55°C for 1 hour. The DNA fragments were then extracted using standard phenol:chloroform:isoamyl alcohol protocol and dissolved in 23 μL of ddH_2_O. Following DNA extraction, the tagmented fragments were PCR amplified using 21 μL of sample, 25 μL NEBNext High-Fidelity 2X PCR Master Mix (New England Biolabs #M0541S) and 2 μL i5 primer and 2 μL i7 primer. The resulting DNA was then cleaned-up and size-selected using 65 μL SPRI beads (Macherey-Nagel NucleoMag #744970). The resulting DNA was resuspended in ddH2O, concentration checked and size distribution visualized using High Sensitivity D1000 ScreenTape (Agilent Technologies #HSD1000) on an Agilent TapeStation 4150 System (Agilent Technologies #G2992AA). Sequencing was done on a NovaSeq X Plus platform, using a paired-end 150 bp per read strategy.

### CUT&Tag bioinformatic analysis

CUT&Tag data were analyzed using Galaxy platform and its public server at *usegalaxy.eu* (Abueg et al., 2024). We followed an analysis pipeline derived from a combination of previously published protocols (Fu et al., 2026; N. Zhang et al., 2025; Zheng et al., 2020). FASTQ files with raw reads were cleaned up and trimmed using Trim Galore (Krueger, 2021). Specifically, we removed the Nextera adapters, any reads with quality cutoff less than 20, and any reads shorter than 30 bp. Quality of cleaned up data was checked and summarized using Falco (de Sena Brandine & Smith, 2021). Cleaned data was then mapped to the built-in TAIR10 genome using Bowtie2 with following parameters: very-sensitive local; no-mixed; no-discordant; fragment length: 10 (-I) < 700 (-X) (Langmead et al., 2009; Langmead & Salzberg, 2012). We then used Filter BAM tool to filter out uninformative reads and remove mitochondrial and chloroplast reads using following conditions: ProperPair: yes, Mapping quality: ≥30, and Reference name of the read: !chrM and !chrPt (Barnett et al., 2011). PCR duplicates were marked/removed using the MarkDuplicates tool and peak calling (H3K27me3) was done using MACS2 callpeak using parameters: Format: - BAMPE; Model: -nomodel; q-value cutoff: 0.05; Broad regions; Broad regions cutoff: 0.1; - SPMR (Feng et al., 2012; Y. Zhang et al., 2008). Files were sorted and consensus peaks identified using bedtools. A region was considered a consensus peak only if a peak had a fold change ≥ 2; q-value < 0.05 and was present in both replicates of a treatment/genotype. Consensus peak files were used as GTF input gene annotation file for featureCounts tool (Liao et al., 2014). Differential peaks were identified using DESeq2 (Love et al., 2014). ChIPseeker was used for peak annotation; with priority set to promoters, 5UTR, 3UTR, exon, intron, downstream and intergenic regions (G. Yu et al., 2015). Gene ontology (GO) enrichment analysis was performed using R package clusterProfiler with *org.At.tair.db* annotation (Yu et al., 2012). Plots were generated using *ggplot2 (Wickham, 2016)*. For visualization of H3K27me3 with H3 background correction, we generated BedGraph files using MACS2 bdgcmp function (Feng et al., 2012; Y. Zhang et al., 2008). BedGraph files were then converted to BigWig file using Convert BedGraph to BigWig tool. Files from replicates were averaged using bigwigAverage (Ramírez et al., 2016) and visualized with Integrative Genomics Viewer (IGV) (Thorvaldsdottir et al., 2013).

### Statistical analyses

Statistically significant differences were assessed by either a Student’s t-test, Fisher’s exact test or one-/two-way ANOVA followed by Tukey’s test using R studio or GraphPad. The used statistical test, number of biological replicates and experimental repeats is indicated in the respective figure legends.

## Supplemental Figures

**Figure S1.**
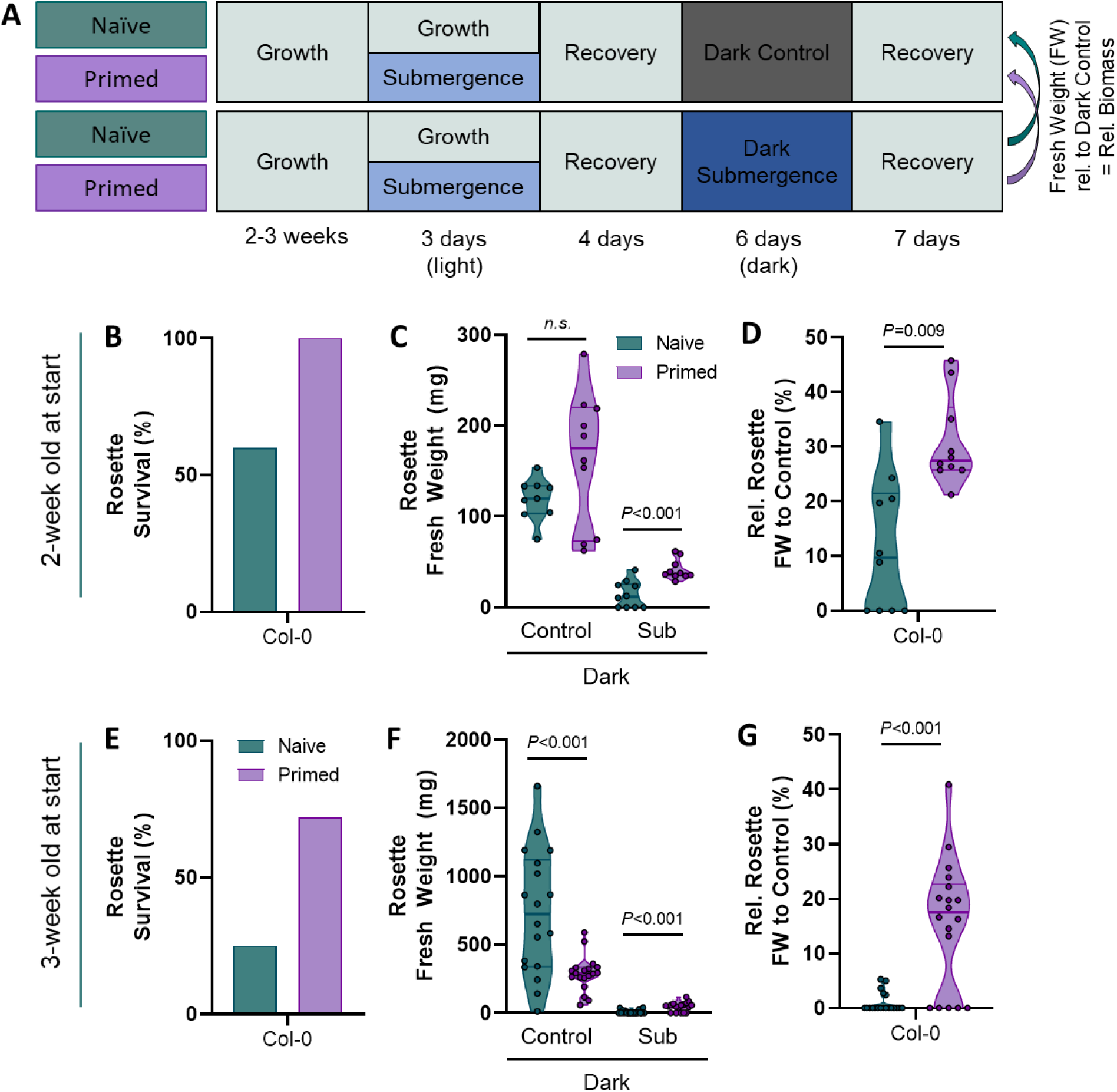
Flooding stress memory is independent of plant age or a priming growth penalty. **(A)** Experimental setup to quantify the effects of light submergence (priming) treatments on subsequent dark- and dark submergence-induced effects on absolute and relative rosette biomass. **(B)** Rosette survival, **(C)** absolute and **(D)** relative to dark control rosette biomass retention of naïve (teal) and submergence primed (violet) of two-week-old Arabidopsis plants at the start of the experimental setup shown in A. **(E)** Rosette survival, **(F)** absolute and **(G)** relative to dark control rosette biomass retention of naïve (teal) and submergence primed (violet) of three-week-old Arabidopsis plants at the start of the experimental setup shown in A. Relative rosette biomass (D, G) was calculated relative to the mean rosette biomass after dark control treatments. P-values are the result of two-tailed Student’s t tests, with n = 10 in B-D (2 experimental repeats with similar results) and n = 18-20 in E-G (1 out of 3 experimental repeats shown with similar results).

**Figure S2.**
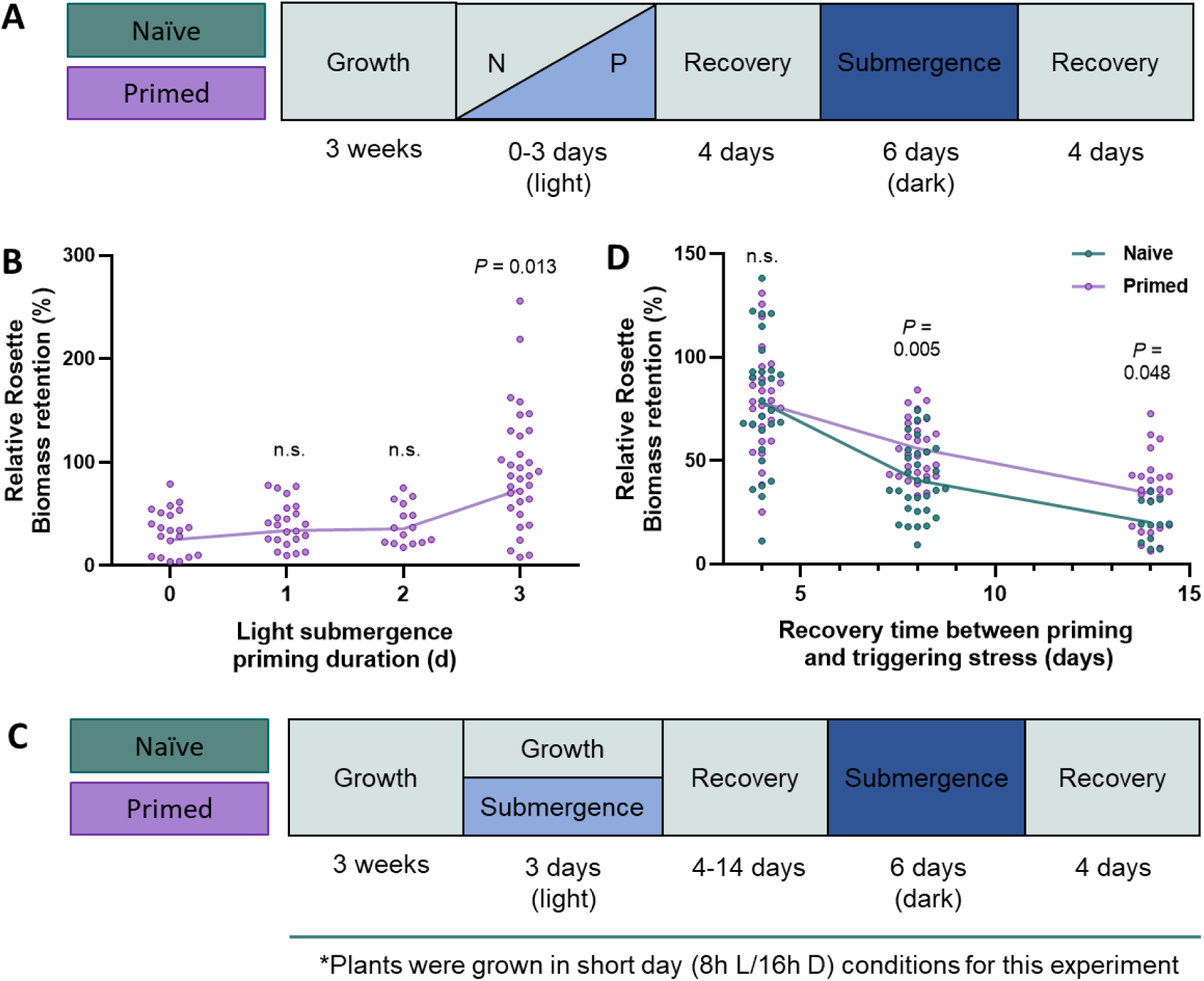
Priming requires 3 days of light submergence and stress memory is maintained for at least 14 days. **(A)** Experimental setup to quantify the effects of light submergence (priming) treatment duration on subsequent dark submergence-induced relative rosette biomass. **(B)** Relative rosette biomass retention of surviving Arabidopsis plants after 0, 1, 2 and 3 days of submergence priming and recovery of 6 days dark submergence. **(C)** Experimental setup to quantify the effects of the recovery duration after light submergence (priming) treatment and subsequent dark submergence. **(D)** Relative shoot fresh weight of primed (violet) and naïve (teal) rosette biomass retention of surviving Arabidopsis plants after 4, 8 and 14 days of recovery of 6-day dark submergence compared to dark controls (Fig. S1). To avoid flowering, plants were grown in short day conditions to allow a longer experimental time, which increased basal flooding tolerance. P-values are the result of a two-tailed Student’s t test comparing naïve (0 days in B) to primed plants, n = 32 (2 experimental repeats).

**Figure S3.**
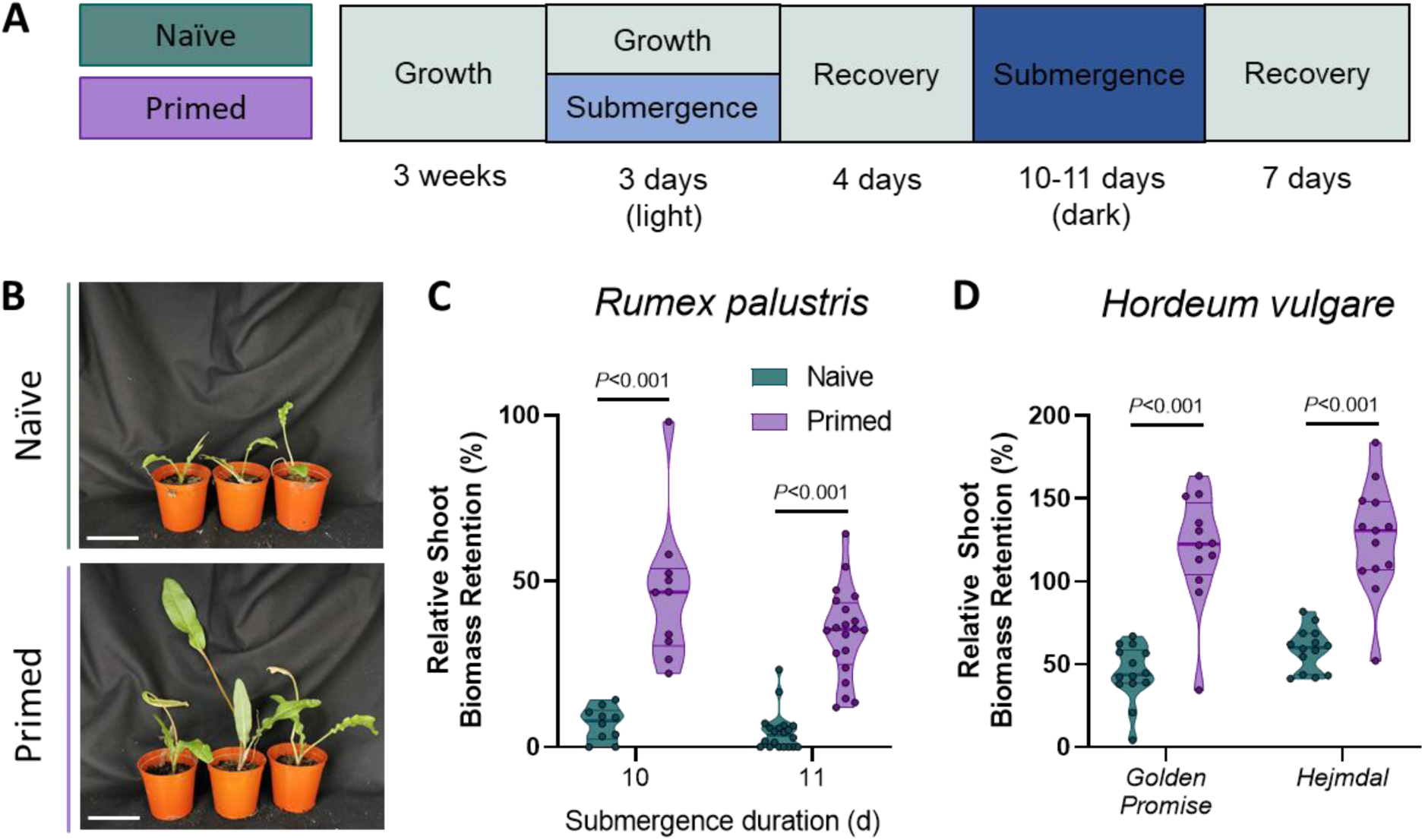
Flooding stress memory is conserved in several angiosperm species. **(A)** Experimental setup to quantify the effects of light submergence priming treatments on dark submergence tolerance. **(B)** Representative images of naïve (teal) and submergence primed (violet) Rumex palustris plants after recovery of 10 days of dark submergence (Scale = 5 cm). Relative shoot fresh weight of primed (violet) and naïve (teal) Rumex palustris **(C)** plants and Hordeum vulgare (barley) genotypes **(D)** after recovery of 10 (for both C and D) or 11-day dark submergence compared to dark controls (Fig. S1). P-values (C and D) are result of two-tailed Student’s t tests, with n = 10 for days and n = 20 for 11 days in C, and n= 12-14 in D.

**Figure S4.**
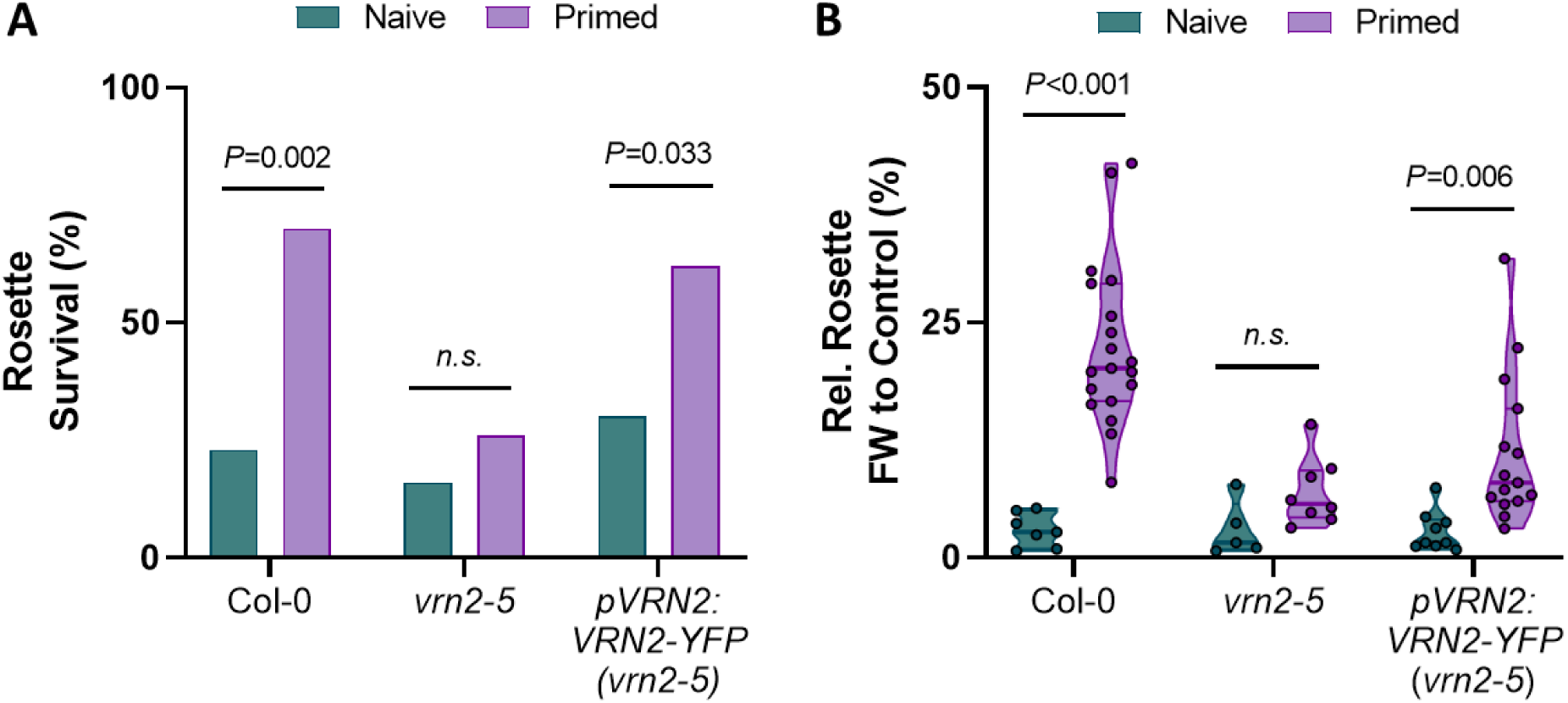
Flooding stress memory is mediated by VRN2. **(A)** Plant survival and **(B)** relative shoot biomass retention of surviving naïve (teal) and submergence primed (violet) Arabidopsis wildtype Col-0, *vrn2-5* mutant and *pVRN2:VRN2-YFP* (*vrn2-5*) complementation plants after recovery of six days of dark submergence. P-values are the result of a chi-squared test in A and a two-tailed Student’s t test in B, with n = 18-20 plants (survived plants plotted only in B). Experiments were repeated three times with similar results.

**Figure S5.**
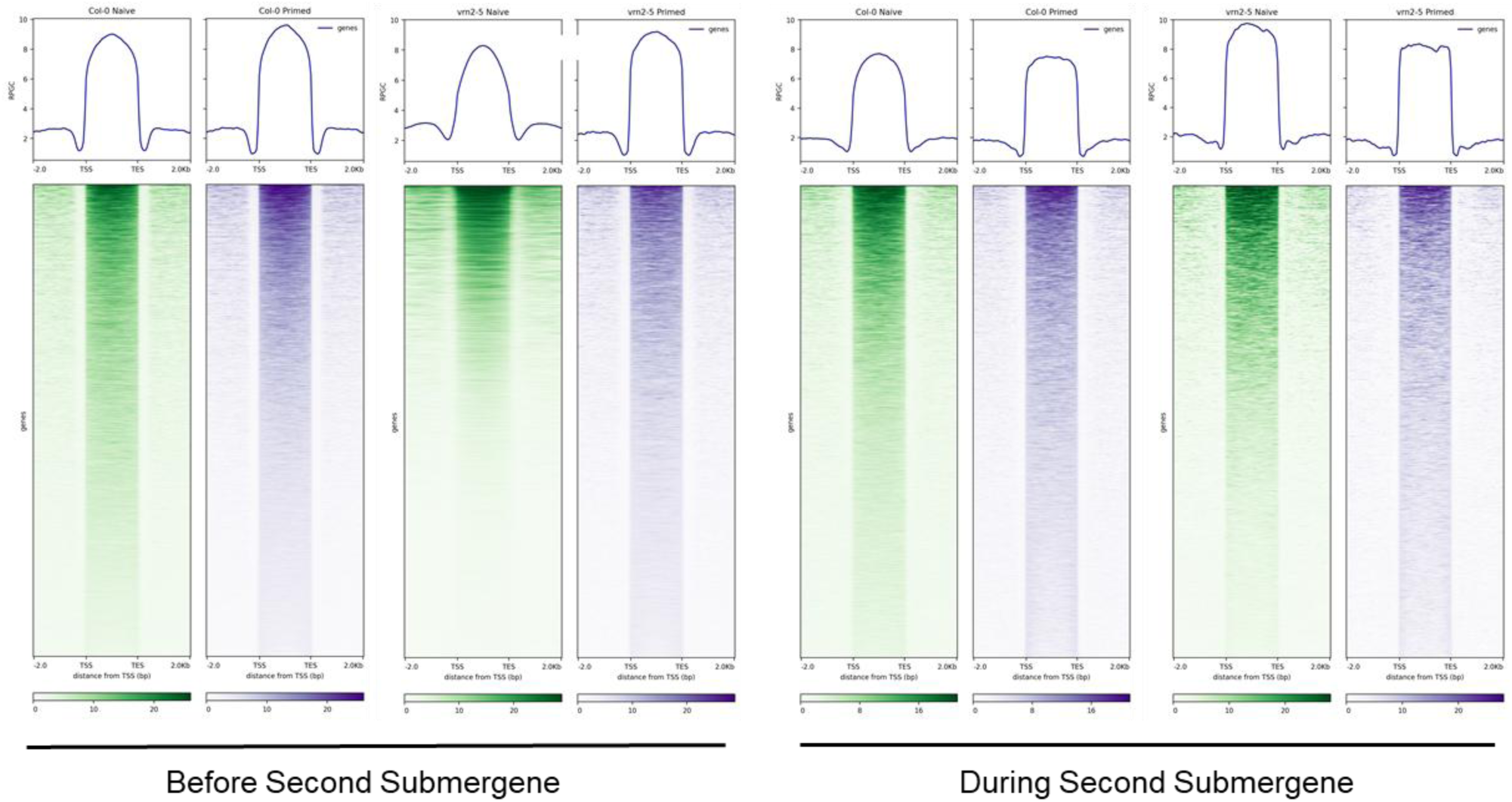
VRN2-dependent global H3K27me3 dynamics during flooding stress memory. Metagene plots (top) and associated heatmaps (bottom) showing the distribution of H3K27me3 / H3 in naïve (green) and primed (purple) Col-0 and *vrn2-5* plants before and during the triggering submergence stress.

**Figure S6.**
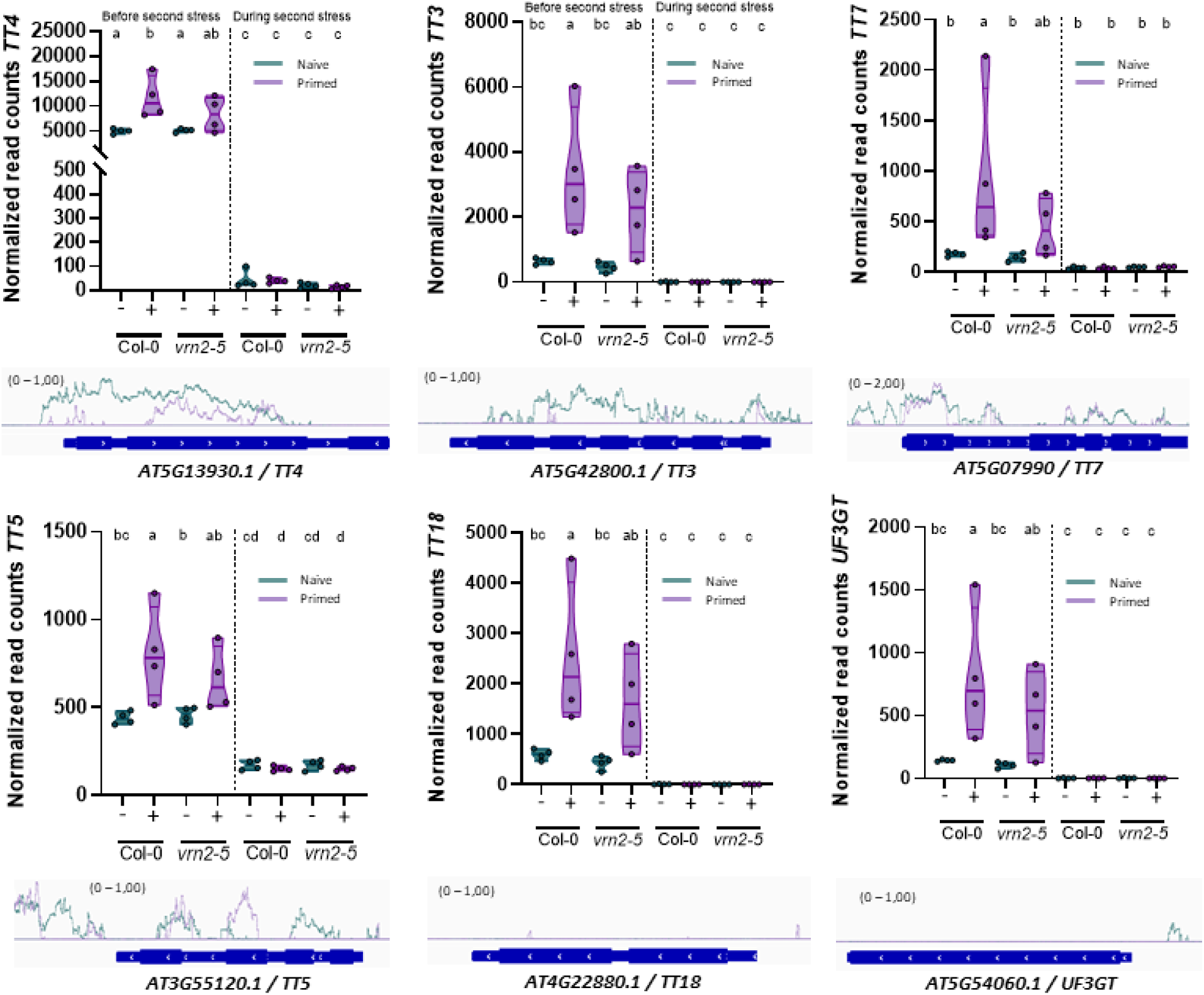
Several anthocyanin biosynthesis genes show H3K27me3 reduction in primed plants, corresponding to its VRN2-dependent type I transcriptional memory pattern. Results extracted from the RNA-Seq data (Figure 2) summarizing the transcript levels of anthocyanin pathway genes in Col-0 and *vrn2-5*, naïve (teal) and primed (violet) plants before and 24h into the dark submergence. Results from four biological replicates are shown as individual datapoints, with lines indicating the median and quartile range. The significance letters are a result of a two-way ANOVA test with Tukey’s HSD (P < 0.05). Coverage plots of H3K27me3 relative to H3 is shown in the indicated naïve (- / teal) and submergence primed (+ / violet) Col-0 rosettes, before the triggering submergence treatment (mean coverage of 2 biological replicates). Data of *TT4* is included for a complete overview of the pathway, but identical to that shown in Fig. 3C & D, and additionally includes transcript levels during the second stress.

**Figure S7.**
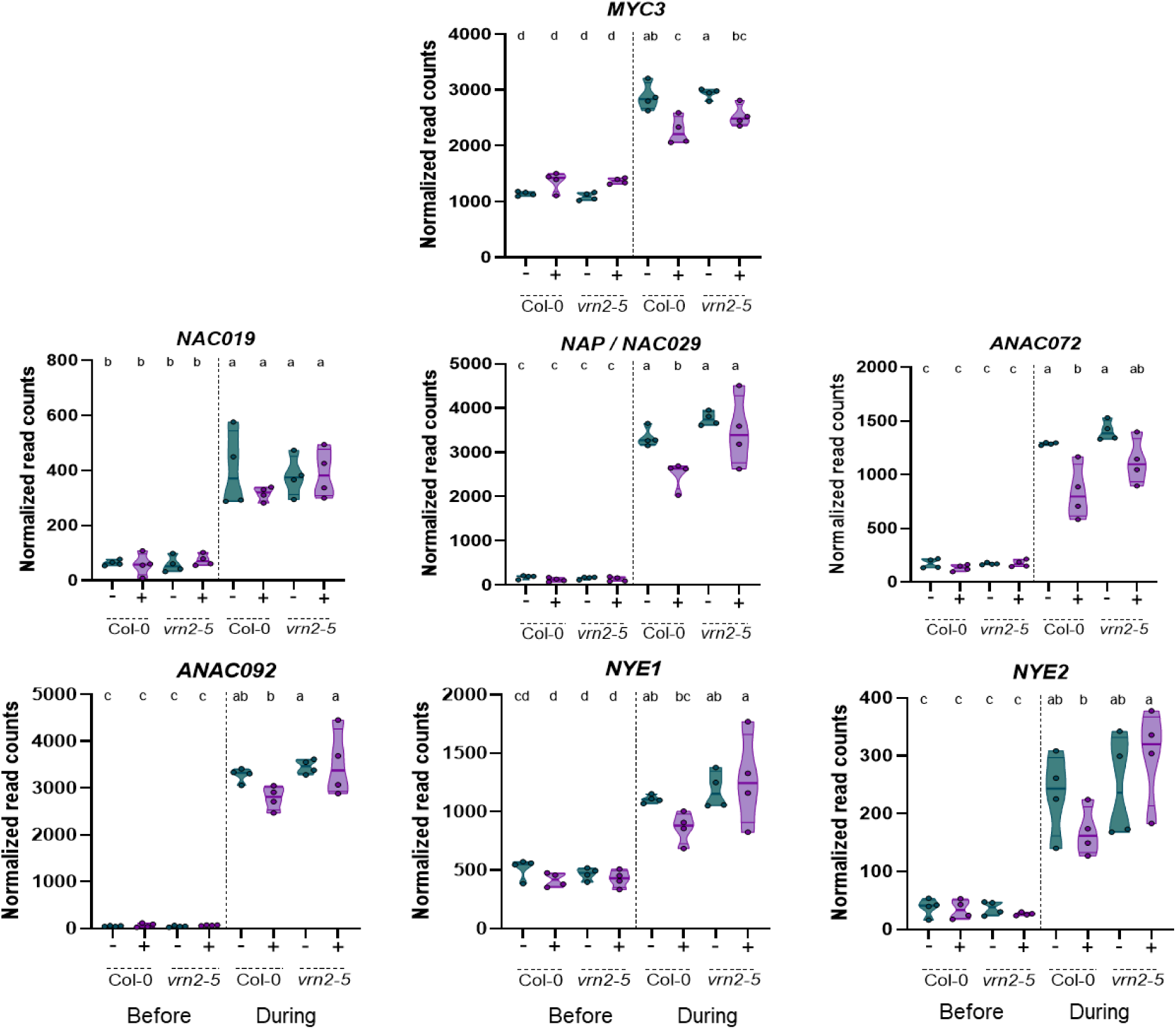
Several senescence-related genes show a VRN2-dependent type II transcriptional memory pattern. Results extracted from the RNA-Seq data (Figure 2, Fig. 4) summarizing the transcript levels of selected senescence-related pathway genes (Fig. 4A, B) in Col-0 and *vrn2-5*, naïve (teal) and primed (violet) plants before and 24h into the dark submergence. Results from 4 biological replicates are shown as individual datapoints, with lines indicating the median and quartile range. The significance letters are a result of a two-way ANOVA test with Tukey’s HSD (P < 0.05).

**Figure S8.**
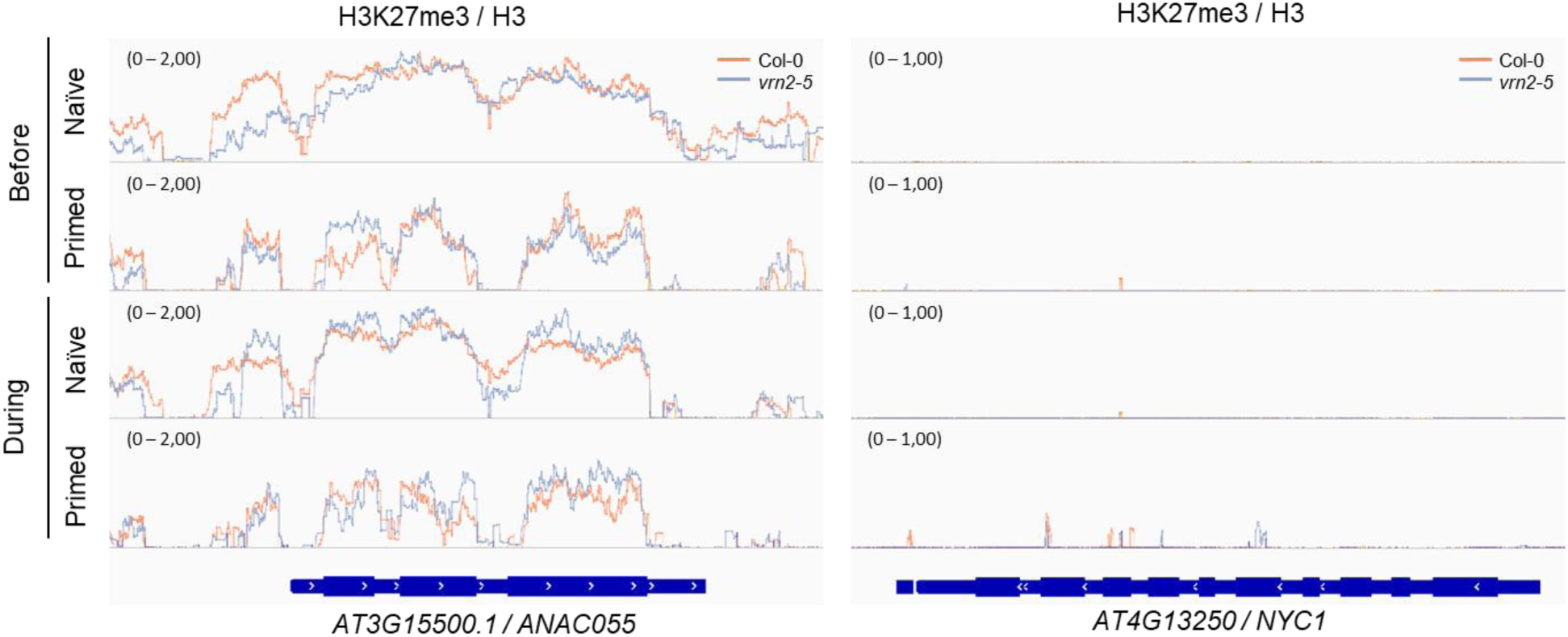
The *NAC055* locus shows VRN2-dependent H3K27me3 enrichment before submergence stress. Coverage plots showing H3K27me3 relative to H3 of the *NAC055* and *NYC1* (involved in senescence regulation) loci in the indicated treated Col-0 wildtype (orange) and *vrn2-5* (blue) plants (mean coverage of 2 biological replicates).

